# The Spectraplakin Short Stop (Shot) Organizes an Acentrosomal Microtubule Network in Early Oogenesis, Essential for Nuclear Positioning

**DOI:** 10.64898/2026.05.13.724832

**Authors:** Fanny Roland-Gosselin, Michel Peroche, Samentha Sahayan Nevil Fernando, Akila Yagoubat, Paul T Conduit, Antoine Guichet, Fred Bernard

## Abstract

Nuclear positioning relies on the coordinated organization of cytoskeletal networks to generate and balance intracellular forces. In the *Drosophila* oocyte, the nucleus undergoes a centering phase during mid-oogenesis before migrating asymmetrically to the cortex in a microtubule (MT)–dependent manner, a process essential for dorso–ventral axis specification. However, the origin and organization of the MT network responsible for nuclear positioning remain to be fully understood.

Here, we identify a previously uncharacterized non-centrosomal Microtubule-Organizing Center (ncMTOC) at the posterior cortex of the oocyte. This ncMTOC is defined by the polarized accumulation of the spectraplakin Short Stop (Shot) together with the MT minus-end–binding protein Patronin prior to nuclear migration. Loss of Shot or Patronin results in posterior displacement of the nucleus, demonstrating that this cortical MT array generates pushing forces that maintain the nucleus at the oocyte center before directional migration. Within this ncMTOC, Shot plays a key role in recruiting both the MT severing enzyme Katanin and the MT polymerase of the XMAP215 family protein Mini-spindle (Msps), ensuring the formation of MTs that emanate from the vicinity of the posterior plasma membrane towards the oocyte cytoplasm. In addition, this ncMTOCs is also associated with γ-TuRC which function independently of centrosomes. Together, these activities establish a dynamic posterior acentrosomal MT network essential for nuclear centering during early oogenesis.

## Introduction

The positioning of the nucleus within a cell is a fundamental aspect of cellular organization and function, influencing key processes such as cell division, polarity establishment, and intracellular transport ^1,2^. In *Drosophila*, proper nuclear positioning during oogenesis is particularly crucial for defining the embryo’s dorsal-ventral polarity ^3,4^. From mid-oogenesis (stages 7/8) onward, the nucleus is positioned in the anterodorsal region of the oocyte, *i.e.*, close to the anterior plasma membrane, adjacent to the nurse cells, and the lateral plasma membrane contacting the follicle cells. The nucleus-associated signaling molecule Gurken is then locally secreted, activating receptors in the adjacent follicle cells, thereby specifying their dorsal fate (González-Reyes et al., 1995; Riechmann and Ephrussi, 2001). Current knowledge is that MTs are the main, if not only, source of forces that displace the nucleus within *Drosophila* oocytes and that proper oocyte nucleus positioning is tightly linked to the organization of a MT network during oogenesis ^7–9^. This functional relationship is reflected in the dynamic changes in nuclear position observed throughout oocyte development.

Before reaching its final location at the anterodorsal corner, nuclei are predominantly located near the anterior membrane at stage 5 and early-stage 6 (stage 6A), whereas at late-stage 6 (stage 6B), they adopt a central position before migrating along two main routes toward either the anterior or the posterior membrane ^10,11^. Previous work indicates that its anterior position at stage 5 is due to an important activity of centrosomes ^11^ that are stereotypically positioned between the nucleus and the posterior plasma membrane of the oocyte. Then a partial decrease of centrosome activity by the MT-associated motor Kinesin I participates to the centration of the nucleus at stage 6B ^11^. However, the mechanisms and forces that maintain the nucleus in a central position, rather than at the posterior, prior to migration, remain to be elucidated.

MTs are dynamic polarized polymers composed of α/β-tubulin heterodimers. MT assembly is initiated at MT organizing centers (MTOCs), which perform three essential functions: MT nucleation, stabilization and anchoring. *In vivo*, MT nucleation relies principally on multi-protein γ-tubulin ring complexes (γ-TuRC). This complex, composed of γ-tubulin and γ-tubulin complex proteins (GCPs), serves as a structural template for the initiation of MT assembly ^12,13^. In addition, recent studies have identified Tumor Overexpressed Gene- (TOG-) domain containing proteins as key regulators that either enhance γ-TuRC-mediated nucleation or trigger MT nucleation independently of γ-TuRC^14–18^. The centrosome is the major MTOC in mitotically dividing cells but differentiated cells commonly use non-centrosomal MTOCs (ncMTOCs) ^14–21^. The relevance of ncMTOCs is particularly significant in the oocyte given that a hallmark of female gametogenesis in most, if not all, species is the loss of centrosomes before fecundation. In *Drosophila* oocytes, centrosome loss occurs during mid-oogenesis from stage 9 onwards ^22,23^; however, before stage 9 centrosomes are active and several types of MTOCs coexist ^7,10,24^. Centrosomes are involved in the positioning of the nucleus ^10,11,25^, while ncMTOCs are at least required for oocyte determination, polarized transport from the nurse cells to the oocyte and nuclear positioning ^14,26–29^. Among the key regulators of ncMTOC organization, the Patronin/CAMSAP family of minus-end-binding proteins ^30^ stabilize MT minus ends by protecting them from depolymerization and by promoting their anchoring at MTOCs. The spectraplakin Short stop (Shot) acts as a cortical MT anchor by linking cortical actin to Patronin/CAMSAP. This interaction has been implicated in the establishment of polarized MT arrays in various cell types, including *Drosophila* oocytes ^27–29^. Another putative MT anchor is Ninein, which has been suggested to stabilize MT minus ends and contribute to ncMT array organization in several cellular systems^14,31–34^. Its role during *Drosophila* oogenesis, however, remains to be fully characterized. Furthermore, the TOG-domain containing protein Mini spindles (Msps), the *Drosophila* orthologue of XMAP215, has been shown to fulfil MT nucleation functions for ncMTOCs, particularly during oocyte determination and, subsequently, for asymmetric transport from nurse cells to the oocyte maintaining its fate ^35,36^.

Here, we have investigated the roles of Shot and Patronin in early nuclear positioning, focusing on the centering step at stage 6, and identified a previously unidentified ncMTOC at the posterior plasma membrane of the oocyte. This structure generates MT-based pushing forces that maintain the nucleus in a central position prior to migration. Shot is a key component of this MTOC, promoting the recruitment of factors involved in MT organization and assembly, including Ninein, Msps and Katanin. Overall, these findings highlight the coordinated and complementary roles of ncMTOC components in regulating nuclear positioning.

## Results

### Loss of Shot Causes Nuclear Positioning Defect in Mid-Oogenesis

Shot and Patronin were previously reported to be enriched along the anterior plasma membrane of the oocyte at stage 10 and excluded from the posterior due to Par-1 activity ^26^. However, using anti-Shot antibodies or a GFP transgene within a BAC (Bacterial Artificial Chromosome), we unexpectedly found that Shot is specifically enriched at the posterior plasma membrane of the oocyte prior to nuclear migration at stages 5 and 6 (Fig. 1A-C). Consistent with previous findings, we also detected its expected enrichment at the anterior plasma membrane from stage 9 onward (data not shown). Interestingly, at stages 5 and 6, Patronin is also enriched at the posterior plasma membrane, in a pattern comparable to that of Shot (Fig. 1D).

**Figure 1.**
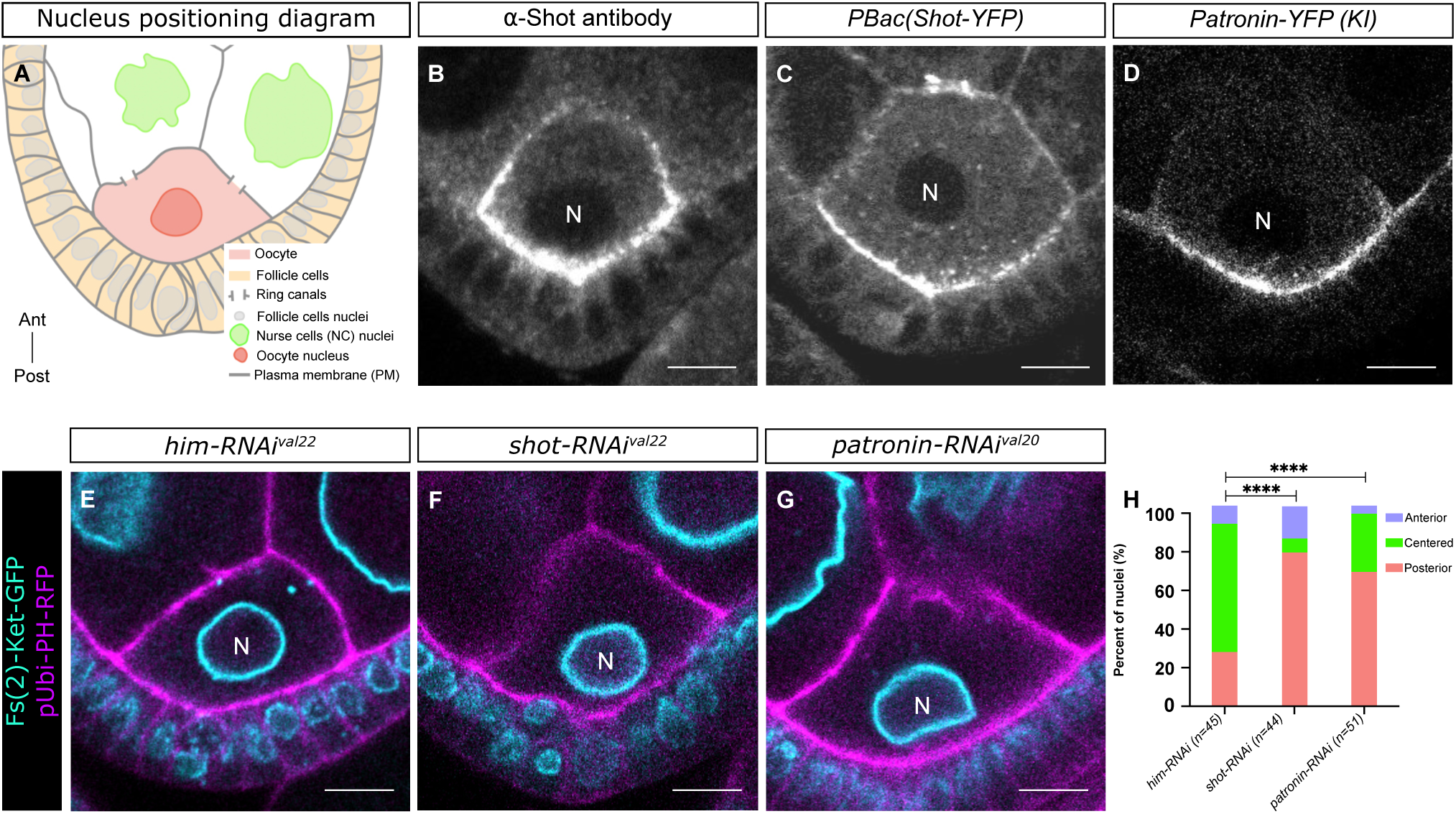
Shot and Patronin are required for nuclear centration. **(A)** Schematic representation of a stage 6B egg chamber. The nucleus (red) is positioned at the center of the oocyte (orange). **(B-D)** Representative images illustrating the localization of Shot and Patronin proteins on stage 6B oocytes, detected using anti-Shot antibody (B), Shot-YFP genomic BAC (Shot-YFP) (C), and Patronin-YFP (CRISPR knock-in) (D). **(E-G)** Representative images illustrating the position of the nucleus in stage 6B oocyte for *him-RNAi^val^*^22^ as control (E), *shot-RNAi^val^*^22^ (F), and *patronin-RNAi^val^*^20^ (G) conditions. The nuclear envelope (cyan) is highlighted by Fs(2)Ket-GFP and the plasma membrane (magenta) by PH-RFP. **(H)** Quantification of the proportions of nuclear positions in the control, *shot-RNAi* and *patronin-RNAi* groups. Positions have been categorized and color-coded as anterior in pale blue, center in green, and posterior in red. n indicates the number of analyzed egg chambers. In all panels, the oocytes are oriented with anterior at the top and posterior at the bottom. Scale bars: 10µm. **** : p<0.0001

To assess the functional relevance of these localization patterns and to determine whether these proteins contribute to MT-associated forces involved in nuclear positioning within the oocyte, we analyzed nucleus positioning in loss-of-function contexts. Because strong *shot* alleles cause defects in oocyte determination and growth^27,29,37^, they are unsuitable for monitoring nuclear positioning; therefore, we opted for RNAi-mediated knockdown. Using the UAS/Gal4 system to drive moderate RNAi expression targeting *shot* with *oskar-Gal4*, we reduced its level while preserving sufficient oocyte size (Fig. S1 E, F). Under these conditions and with two different RNAi lines, we observed that at stage 6B, the oocyte nucleus was predominantly positioned posteriorly compared to control conditions, where nuclei remained centered (Fig. 1 E-F, H and Fig. S1 B-D). Similarly, using *matTub-Gal4* to drive *patronin-RNAi*, we observed a posterior shift of the oocyte nucleus (Fig. 1 G-H). Taken together with their observed localizations, these findings support the hypothesis that Shot and Patronin contribute to organizing MTs required to push the nucleus away from the posterior cortex toward the center of the oocyte.

### Loss of Shot Causes MTs Defect in Mid-Oogenesis

To investigate whether Shot and Patronin regulate nuclear positioning via MTs in stage 6 oocytes, we examined the global MT network in live egg chambers using SiR-tubulin, a fluorogenic, cell-permeable, highly specific MT probe. Oocyte MT organization was further analyzed using a *ubi-PH-RFP* transgene to mark plasma membranes and outline the oocyte contour, thereby providing a reference for spatial organization (Fig. 2A). A significant decrease in the SiR-tubulin signal was observed in both the *shot-RNAi* and *patronin-RNAi* oocytes (Fig. 2B–E), including in the region of the posterior Plasma Membrane (pPM) (Fig. 2E). This indicates that both proteins are important for forming a dense MT network in oocytes, including close to the pPM. We next asked whether these proteins could organize a MT network from the posterior plasma membrane. To test this hypothesis, we recorded time-lapse movies of live oocytes expressing endogenously GFP-tagged End-binding protein 1 (EB1-GFP). EB1 is a MT associated protein that binds to polymerizing MT plus ends. Thus, tracking EB1 signals allows visualization of MT plus-end dynamics and, consequently, determination of MT polarity. In stage 6 oocytes, EB1 signals were stronger near the posterior plasma membrane, reflecting higher MT density (Fig. 2F, arrowheads). Frame-by-frame analysis of the movies revealed comets emerging from the pPM and moving towards the center of the oocyte, hereafter referred to as posterior Plasma Membrane Arising Comets (pPMACs). These pPMACs support the existence of a MT network emanating from the pPM (Fig. 2G). To globally assess and quantify the network we developed a semi-automated image segmentation and tracking method to vectorize comet trajectories (see Material and Methods) (Fig. 2H). Representing tracks as vectors allowed us to generate mathematical objects in which the direction corresponds to the orientation of MT growth, and the magnitude corresponds to comet length over time (Fig. 2H”). This method allowed us to assess growth directionality, polymerization speed, and spatial organization. Applying this analysis to movies of oocytes from control, *shot-RNAi* and *patronin-RNAi* conditions revealed a significant decrease in the number of MTs growing from the posterior plasma membrane in both *shot-RNAi* and *patronin-RNAi* compared to controls (Fig. 2I). Interestingly, the speed of the remaining pPMACs in both *shot-RNAi* and *patronin-RNAi* oocytes was unchanged, suggesting that Shot and Patronin control pPMAC number, rather than their dynamics (Fig. 2J). These findings support a role for Shot and Patronin in MT organization and indicate that, at stage 6 of *Drosophila* oogenesis, the posterior plasma membrane of the oocyte functions as a ncMTOC.

**Figure 2.**
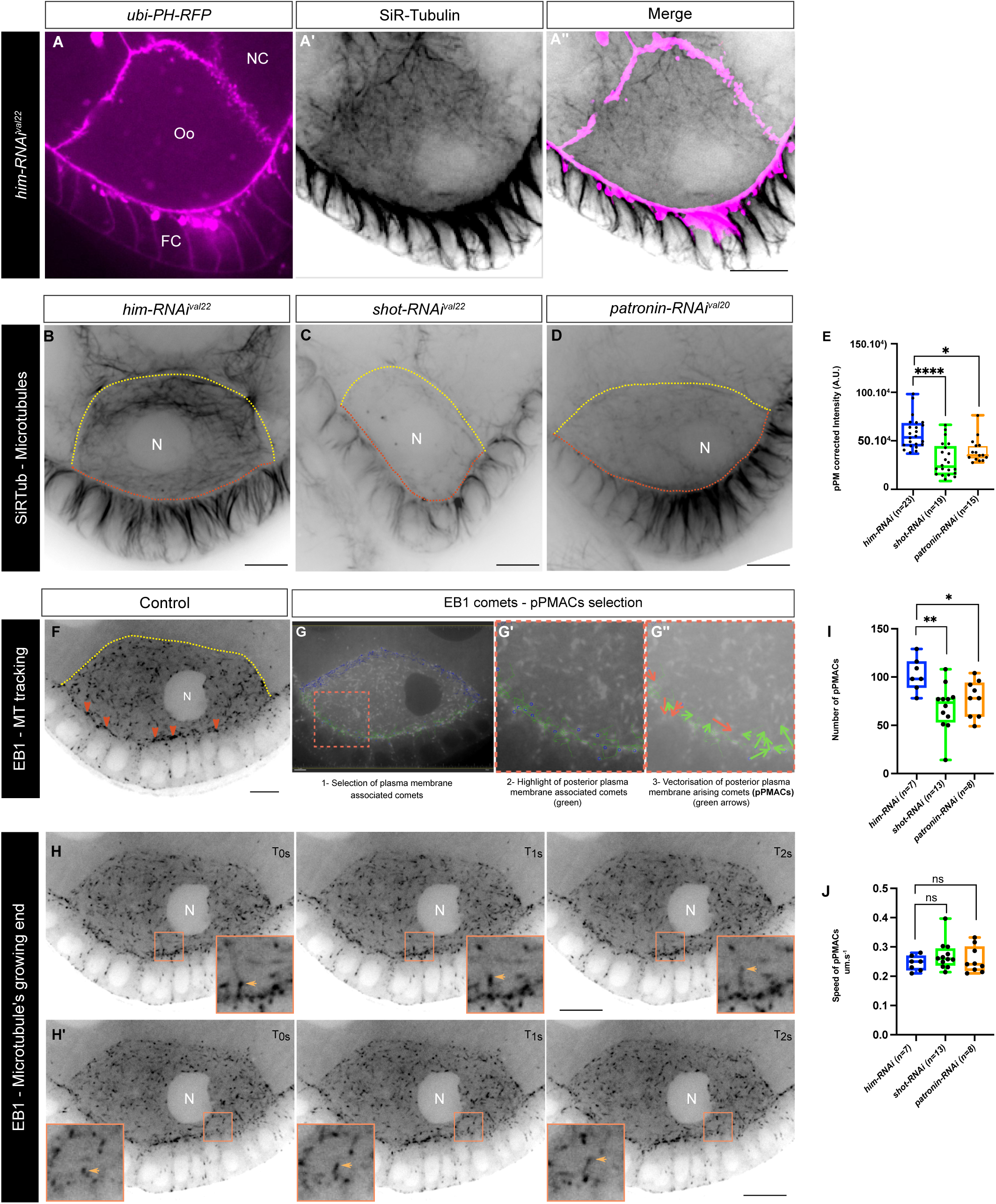
Shot and Patronin are essential for MT network integrity. **(A-D)** Stage 6B egg chambers stained with the SiR-Tubulin probe that reveals MT networks. Control oocyte (*osk-Gal4, ubi-PH-RFP; UAS-him-RNAi^val^*^22^) (A-A’’) delimited by the signal of the *ubi-PH-RFP* (A) exhibits a dense network of MTs (A’); merge in A’’. Representative examples are shown for *him-RNAi^val^*^22^ as control (B), *shot-RNAi^val^*^22^ (C), and *patronin-RNAi^val^*^20^ (D) conditions. The yellow and orange dashed lines, placed using ubi-PH-RFP, highlight the anterior and posterior plasma membranes of the oocyte, respectively. **(E)** Quantification of MT corrected intensity at the vicinity of the posterior plasma membrane (pPM) of oocytes in the control, *shot-RNAi^val^*^22^ and *patronin-RNAi^val^*^20^ conditions. **(F)** MT plus end extremities were revealed by EB1::SSS::GFP in stage 6B oocytes. EB1 comets (red arrowheads) emerged from the posterior plasma membrane. **(G-G’’)** The Tracking module of IMARIS software identifies comet trajectories based on displacement of the EB1::SSS::GFP signal (G). Tracks that include at least one time point within the 10-pixel-wide posterior ROI are classified as posterior trajectories (green in G′). Trajectory vectorization allows determination of the angle of each trajectory relative to the horizontal axis. Comets emerging from the posterior plasma membrane (green arrows) exhibit negative angles and are categorized as pPMACs, whereas comets directed toward the posterior plasma membrane (red arrows) display positive angles (G’). **(H, H’)** Independent time series of MT plus end movement from the posterior cortex (insets and orange arrowhead). **(I)** Quantification of pPMACs in control (*him-RNAi^val^*^22^), *shot-RNAi^val^*^22^ and *patronin-RNAi^val^*^20^ conditions. **(J)** Quantification of pPMAC speed in control (*him-RNAi^val^*^22^), *shot-RNAi^val^*^22^ and *patronin-RNAi^val^*^20^ conditions. In all panels, the oocytes are oriented with anterior at the top and posterior at the bottom. Scale bars: 10µm. * : p<0.05 ; ** : p<0.01 ; **** : p<0.0001. N: oocyte nucleus; FC: follicle cells, Oo: oocyte.

### γ-Tubulin independently of centrosome Causes Nuclear Positioning and MTs Defect in Mid-Oogenesis

We investigated the localization of γ-tubulin in order to identify how microtubules, which are required to push the nucleus away from the posterior cortex, are nucleated. In *Drosophila*, γ-Tubulin is encoded by two genes: γ*-Tub37C* (γ*-Tubulin at 37C*) and γ*-Tub23C*(γ*-Tubulin at 23C*). Analysis of their localization in stage 6B egg chambers revealed that endogenously tagged γ-Tub23C-GFP was specifically detectable in follicle cells at the level of the centrosome, whereas barely any signal was detected in the oocyte (Fig. 3A). By contrast, endogenously tagged γ*-Tub37C-mCherry*, which is germline-specific, exhibited a diffuse cytoplasmic signal together with strong enrichment at centrosomes (Fig. 3B). RNAi mediated depletion of γ*-Tub37C* resulted in posterior displacement of the oocyte nucleus at stage 6B, compared to control conditions in which nuclei remained predominantly centered (Fig. 3C) indicating that γ-Tubulin is required for nuclear centration at that stage. Monitoring MT distribution using the SiR-Tubulin probe following RNAi-mediated depletion of *γ-Tub37C* reveals a significant reduction in MT density, as reflected by decreased SiR-Tubulin signal intensity (Fig. 3D, E), suggesting a role for γ-Tub37C in MT organization in stage 6B oocytes. We then assessed its function at the posterior plasma membrane by tracking EB1 to visualize MT plus-end dynamics and quantifying pPMAC number and speed. While pPMAC number was unchanged upon γ*-Tub37C* knockdown, their velocity was reduced (Fig. 3F, G). This contrasts with Shot and Patronin depletions, which specifically decrease pPMAC numbers, without affecting their dynamics (Fig. 2I-J), suggesting that γ-Tub37C and Shot/Patronin act through distinct pathways.

**Figure 3.**
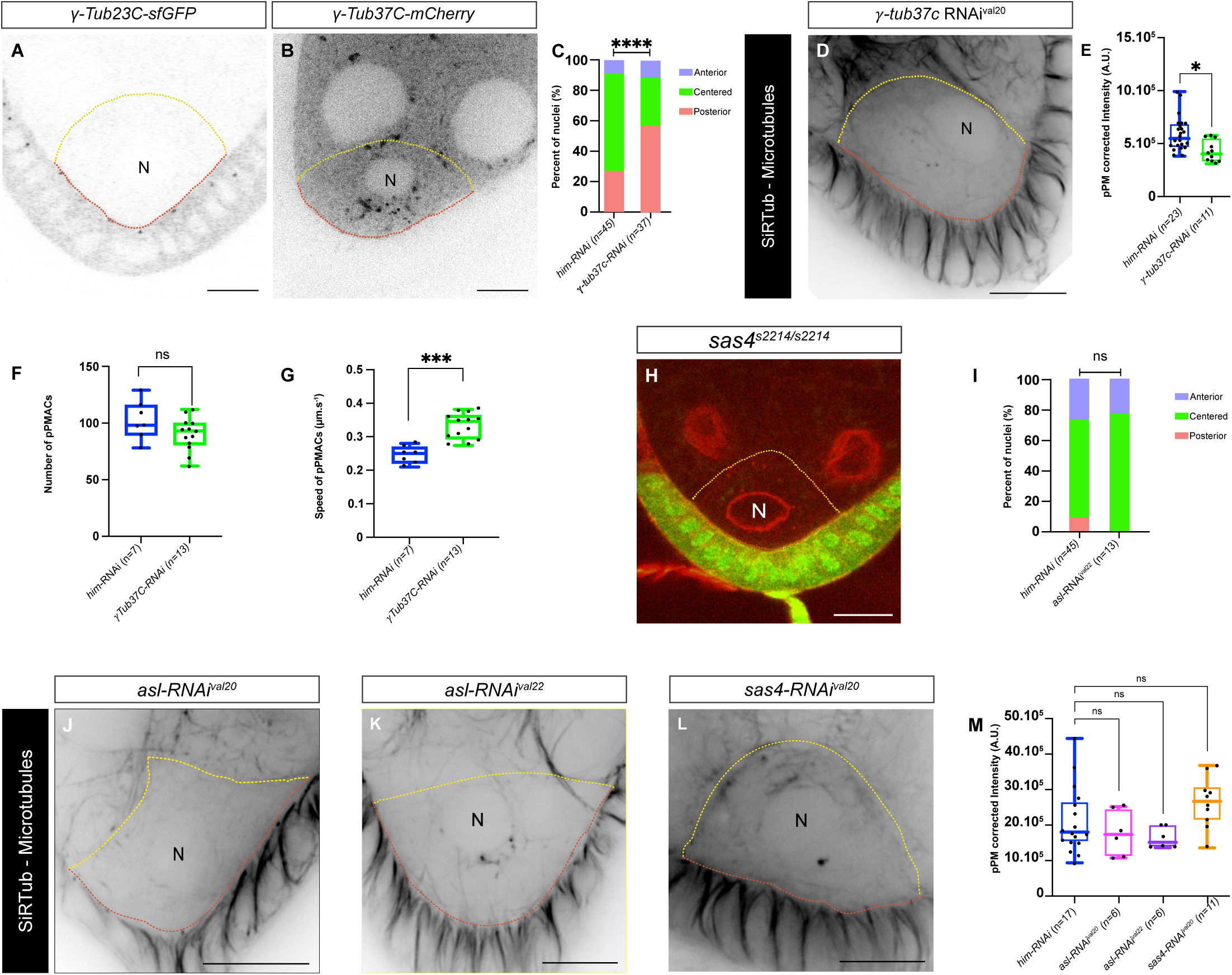
The γ-Tub37C contributes to the organization of the MT network at the pPM. **(A)** Representative fluorescent image of an oocyte expressing endogenously tagged γ*-Tubulin 23C-GFP*. Note that no signal is detected in the oocyte, whereas signals corresponding to the centrosomes are detected in the overlying follicle cells. **(B)** Representative fluorescent image of an oocyte expressing endogenously tagged γ*-tubulin 37C-GFP*. The yellow and orange dashed lines, placed using ubi-PH-RFP, highlight the anterior and posterior plasma membranes of the oocyte, respectively. **(C)** Quantification of the proportions of nuclear positions in the control (*him-RNAi^val^*^22^) and γ*-tubulin 37C-RNAi^val20^*via RNAi transgenes. Positions have been categorized and color-coded as anterior in pale blue, center in green, and posterior in red. n indicates the number of analyzed egg chambers. **(D)** Representative image of Stage 6B egg chamber labeled with the SiR-Tubulin probe to reveal MT with γ*-tubulin 37C-RNAi^val^*^20^ condition. Dotted lines outline the oocyte contour; the anterior cortex is shown in yellow and the posterior cortex in orange. **(E)** Quantification of MT corrected intensity at the vicinity of the posterior plasma membrane (pPM) of oocytes in the control and γ*-tubulin 37C-RNAi^val^*^20^ conditions. **(F, G)** Quantification of pPMACs numbers (I) and speed (J) in control and γ*-tubulin 37C-RNAi^val^*^20^ conditions. **(H)** Homozygous *Sas4^s^*^2214^ mutant clones marked by the absence of nuclear GFP (Green) exhibit a nucleus in a central position. The yellow dashed line indicates the contour of the oocyte. **(I)** Quantification of the proportions of nuclear positions in the control (*him-RNAi^val^*^22^) and *asterless-RNAi^val^*^22^ groups. Positions have been categorized and color-coded as anterior in pale blue, center in green, and posterior in red. n indicates the number of analyzed egg chambers. **(J-L)** Representative images of Stage 6B egg chambers labeled with the SiR-Tubulin probe to reveal MT in *asl-RNAi^val^*^20^ (J), *asl-RNAi^val^*^22^ (K) and *Sas4-RNAi^val^*^20^ (L) conditions. *asl-RNAi^val^*^20^ and *asl-RNAi^val^*^22^ are two independent RNAi lines. The yellow and orange dashed lines, placed using ubi-PH-RFP, highlight the anterior and posterior plasma membranes of the oocyte, respectively. **(M)** Quantification of MT corrected intensity at the vicinity of the posterior plasma membrane (pPM) of oocytes in the control, *asl-RNAi^val^*^20^, *asl-RNAi^val^*^22^ and *Sas4-RNAi^val^*^20^ conditions. Dotted lines outline the oocyte contour; the anterior cortex is shown in yellow and the posterior cortex in orange. Scale bars: 10µm. * : p<0.05 ; ** : p<0.01 ; *** : p<0.001 ; **** : p<0.0001; ns, not significant. N: oocyte nucleus.

We previously showed that centrosomes contribute to nuclear migration, as in their absence the nucleus follows a slower trajectory and predominantly migrates the posterior plasma membrane ^10^, although no defects in antero-dorsal positioning were observed at stage 7-8. ^10^In the light of nuclear positioning defects observed upon γ-Tub37C depletion, we next examined the effects of depleting the centrosome-associated proteins Asterless (Asl) and Spindle assembly abnormal 4 (Sas-4) on nuclear positioning at stage 6B. In contrast to γ-Tub37C depletion, RNAi-mediated knockdown of *asl* did not result in any significant change in nuclear position compared to control (Fig. 3I). Consistently, the nucleus remained properly centered in germline clones carrying the Sas4^S2214^ loss-of-function allele (Fig. 3H). In agreement with these observations, MT network organization was unaffected upon RNAi-mediated depletion of either *asl* or *sas-4* (Fig. 3J-M).

Together, these results indicate that, despite not being specifically enriched at the pPM compared to Shot, γ-Tub37C contributes to the organization of the posterior MT network that centers the nucleus at stage 6B, independently of its canonical centrosomal functions.

### Katanin promotes MTs in the posterior cortex

In order to further understand the process by which Shot organize the MTs, we therefore performed a candidate screen to identify known MT regulators that are enriched at the posterior cortex of stage 6 oocytes. We identified Ninein (Fig. 4A), Katanin (Fig 4D) and Mini-spindles (Msps) (see below, Fig. 5).

**Figure 4.**
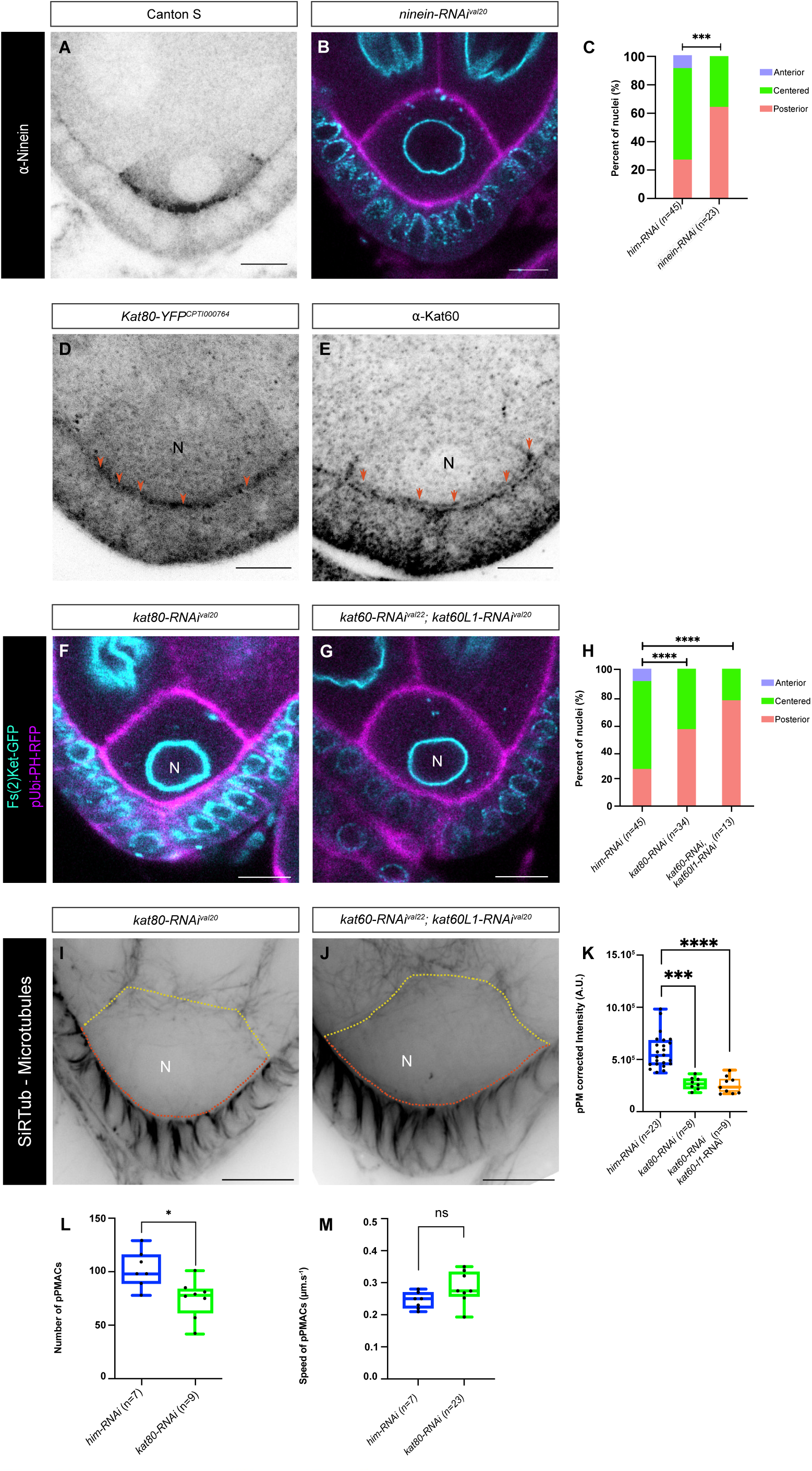
Katanin is required for MT network integrity and nuclear centration. **(A)** Representative image of wild-type stage 6B oocyte labeled Ninein antibody. **(B)** Representative images illustrating the position of the nucleus in stage 6B oocyte expressing *ninein-RNAi^val^*^20^. The nuclear envelope (cyan) is highlighted by Fs(2)Ket-GFP and the plasma membrane (magenta) by PH-RFP. **(C)** Quantification of the proportions of nuclear positions in the control and *ninein-RNAi^val^*^20^ groups. Positions have been categorized and color-coded as anterior in pale blue, center in green, and posterior in red. n indicates the number of analyzed egg chambers. **(D, E)** Representative images of wild-type stage 6B oocyte expressing Kat80-YFP genomic bac (D) and labeled with Kat60 antibody (E). The red arrowheads illustrate the association with the plasma membrane. **(F-H)** Representative images illustrating the position of the nucleus in stage 6B oocyte for *kat80-RNAi^val^*^20^ (F), and joint inactivation of *Kat60* and *Kat60L1* via RNAi (G) conditions and quantifications (H). Positions have been categorized and color-coded as anterior in pale blue, center in green, and posterior in red. n indicates the number of analyzed egg chambers. **(I-K)** Representative images of Stage 6B egg chambers labeled with the SiR-Tubulin probe to reveal MT with *kat80-RNAi^val^*^20^ (I), and *kat60-RNAi^val^*^22^*; kat60L1-RNAi^val^*^20^ (J) conditions. The yellow and orange dashed lines, placed using ubi-PH-RFP, highlight the anterior and posterior plasma membranes of the oocyte, respectively. Quantification of MT corrected intensity at the vicinity of the posterior plasma membrane (pPM) of oocytes in the control, *Kat80-RNAi* and joint inactivation of *Kat60* and *Kat60L1* via RNAi conditions (K). **(L-M)** Quantification of pPMACs numbers (L) and speed (M) in control and *kat80-RNAi^val^*^20^ conditions. In all panels, the oocytes are oriented with anterior at the top and posterior at the bottom. Scale bars: 10µm. * : p<0.05 ; ** : p<0.01 ; *** : p<0.001 ; **** : p<0.0001; ns, not significant. N: oocyte nucleus

**Figure 5.**
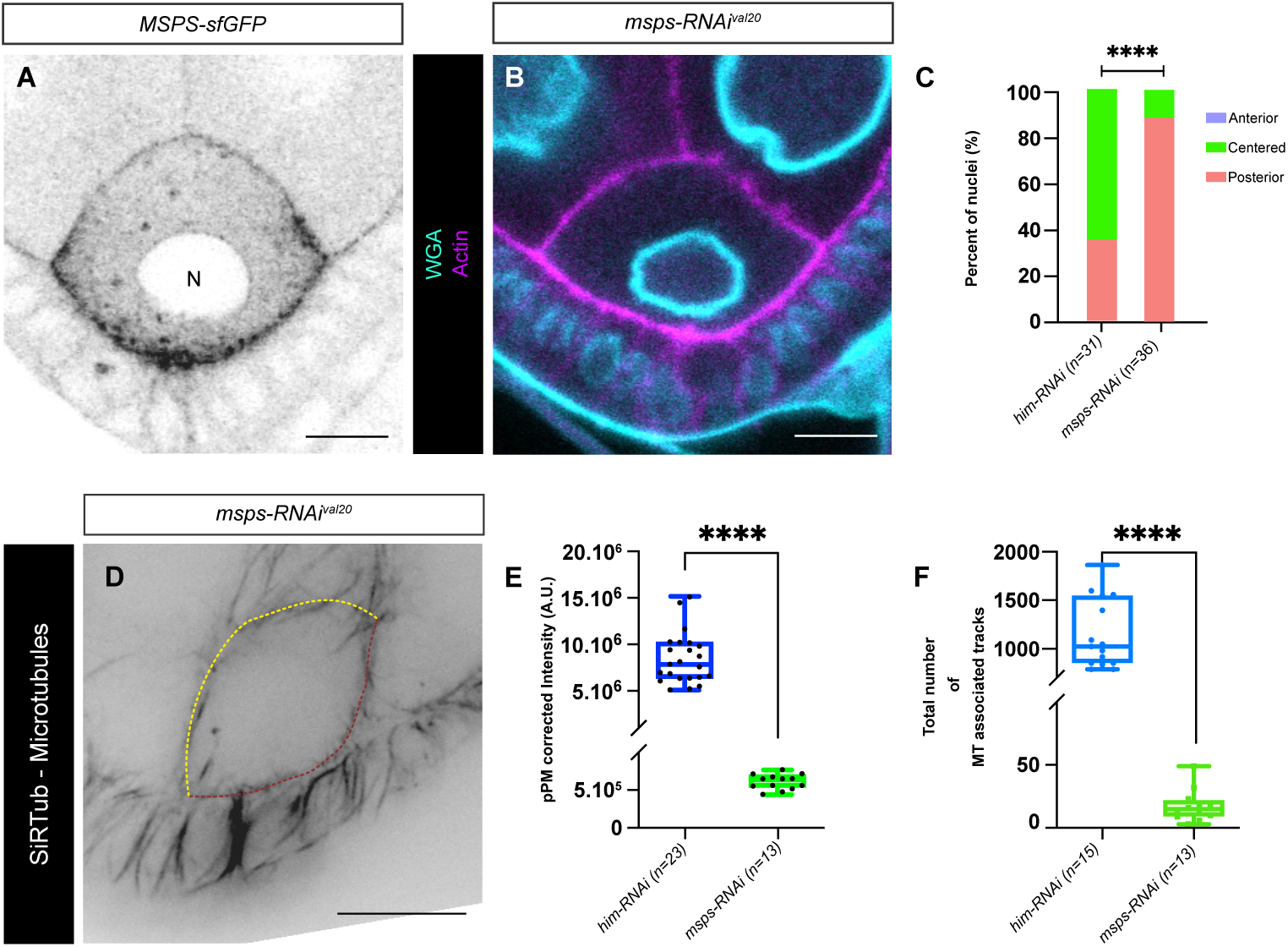
Msps contributes to the Shot defined ncMTOC. **(A)** Representative fluorescent image of an oocyte expressing endogenously tagged Msps-GFP. **(B)** Representative images illustrating the position of the nucleus in stage 6B oocyte for *msps-RNAi* conditions. The nuclear envelope (cyan) is highlighted by WGA probe and the oocyte contour (magenta) by phalloidin. **(C)** Quantification of the proportions of nuclear positions in the control (*him-RNAi^val^*^22^) and *msps-RNAi^val^*^20^. Positions have been categorized and color-coded as anterior in pale blue, center in green, and posterior in red. n indicates the number of analyzed egg chambers. **(D)** Representative images of Stage 6B egg chambers labeled with the SiR-Tubulin probe to reveal MT with *msps-RNAi^val^*^20^ conditions. The yellow and orange dashed lines, placed using ubi-PH-RFP, highlight the anterior and posterior plasma membranes of the oocyte, respectively. **(E)** Quantification of MT corrected intensity at the vicinity of the posterior plasma membrane (pPM) of oocytes in the control (*him-RNAi^val^*^22^) and *msps-RNAi^val^*^20^ conditions. **(F)** Quantification of MT-associated track numbers in the control (*him-RNAi^val^*^22^) and *msps-RNAi^val^*^20^ conditions. In all panels, the oocytes are oriented with anterior at the top and posterior at the bottom. Scale bars: 10µm. **** : p<0.0001; N: oocyte nucleus

Ninein, is a conserved microtubule minus-end–associated protein implicated in the organization and anchoring of non-centrosomal microtubule arrays. In *Drosophila*, it has been shown to localize to microtubule-organizing sites in several tissues, where it contributes to stabilizing microtubule minus ends and supporting cellular polarity^14,31–34^. Its depletion using *ninein-RNAi* led to defects in nuclear positioning at stage 6B, where nuclei significantly localized toward the posterior membrane of the oocyte (Fig. 4B-C).

Katanin belongs the AAA+ ATPases family and forms an octamer consisting of a regulatory subunit, Katanin 80 and catalytic subunits Katanin 60 proteins ^38–41^. In *Drosophila*, the Katanin 60 proteins are encoded by two genes, *Katanin 60* (*Kat60*) and *Katanin p60-like 1* (*Kat60-L1*) ^42,43^. Using anti-Kat60 antibodies and Kat80-GFP protein-trap line, we observed that both Kat60 and Kat80 were enriched at the posterior cortex of stage 6B oocytes (Fig. 4D, E). Moreover, RNAi-mediated co-depletion of *Kat60* and *Kat60-L1* or single depletion of Kat80 resulted in a higher frequency of posteriorly positioned nuclei (Fig. 4 F-H). Moreover, quantification of MT density with the SiR-tubulin probe revealed a significant reduction in *Kat60/Kat60-L1 RNAi* oocytes (Fig. 4I-K). These phenotypes mirror those seen in *Shot-RNAi* and *patronin-RNAi* oocytes, suggesting that Katanin activity contributes to the MT network.

We then examined Katanin’s role in organizing MTs at the posterior plasma membrane. To do so, we tracked EB1 signals to allows visualization of MT plus-end dynamics and monitored the number of pPMACs in *Kat80-RNAi* compared to *him-RNAi* controls. We observed a significant decrease in pPMAC number upon *Kat80* knockdown, whereas their properties, such as speed, remained unchanged (Fig. 4L-M). This indicates that Katanin contributes specifically to the formation of MTs emanating from the pPM, suggesting that Katanin-mediated MT severing promotes the generation of new MTs at the posterior cortex of stage 6 oocytes.

### Msps promotes new MTs in the posterior cortex

Our screen for posteriorly enriched MT-associated proteins, also identified the TOG-domain proteins Msps. Using a CRiSPR line expressing GFP-tagged Msps, we found that Msps is present in the oocyte, as previously reported ^35,36^ but is also particularly enriched at the posterior plasma membrane of stage 6 oocyte (Fig. 5A). Msps is the *Drosophila* ortholog of XMAP215/TOG family proteins. It exhibits MT nucleation properties and has been reported to play a fundamental role in MT dynamics and mitotic spindle assembly ^44^. RNAi-mediated knockdown of *msps* induces growth defects (Fig. S1 G), consistent with its critical role in MT organization within the oocyte, as previously reported ^36^.

To maintain relatively normal oocyte size while efficiently depleting *msps*, we induced RNAi against *msps* using the *osk-Gal4* driver. Under these conditions, most nuclei were positioned at the posterior (Fig. 5B-C). We then assessed MT density in the oocyte following this milder RNAi-mediated depletion of *msps* using SiR-Tubulin and found a significant reduction compared to *him-RNAi* controls (Fig. 5 D-E). To examine the specific role of Msps in organizating the posterior MT network, we tracked EB1 signals to visualize MT plus-end dynamics and assess MT polarity. EB1-positive comet numbers were strongly reduced compared to controls; thus, we quantified total comet number which was significantly decreased (Fig. 5F). Given that reduced Msps activity affects nuclear positioning and its enrichment at the posterior in stage 6 oocytes, these results suggest that Msps contributes to the formation of MTs emanating from the posterior plasma membrane.

### Shot acts as a pPM hub for ncMTOC localization

The posterior localization of an ncMTOC and the enrichment of MT-associated proteins at the posterior of the oocyte raise the question of the underlying cue that directs these spatial distributions. Shot is a central component of multiple ncMTOCs, and its localization correlates with sites of active MT polymerization, for example, it is enriched at the anterior at stage 10, where polymerization activity is intense ^26^. We therefore tested whether Shot is required for the localization of the MT-associated proteins we identified. To this end, we examined their distribution in Shot-depleted oocytes. Under these conditions, Ninein localization is strongly disrupted, with a marked reduction of its accumulation at the posterior (Fig. 6A-C). Kat60 association with the posterior plasma membrane is also strongly reduced upon Shot knockdown (Fig. 6D-F). Finally, Msps association with the posterior membrane is greatly diminished and accompanied by a diffuse redistribution throughout the cytoplasm (Fig. 6G–I). These results indicate that Shot is required to recruit proteins to the posterior region that promote MT assembly, Msps and Katanin, as well as proteins that anchor and stabilize MTs, namely Ninein and Patronin. Together, this suggests that Shot likely functions as a central scaffold that organizes and stabilizes the posterior ncMTOC.

**Figure 6.**
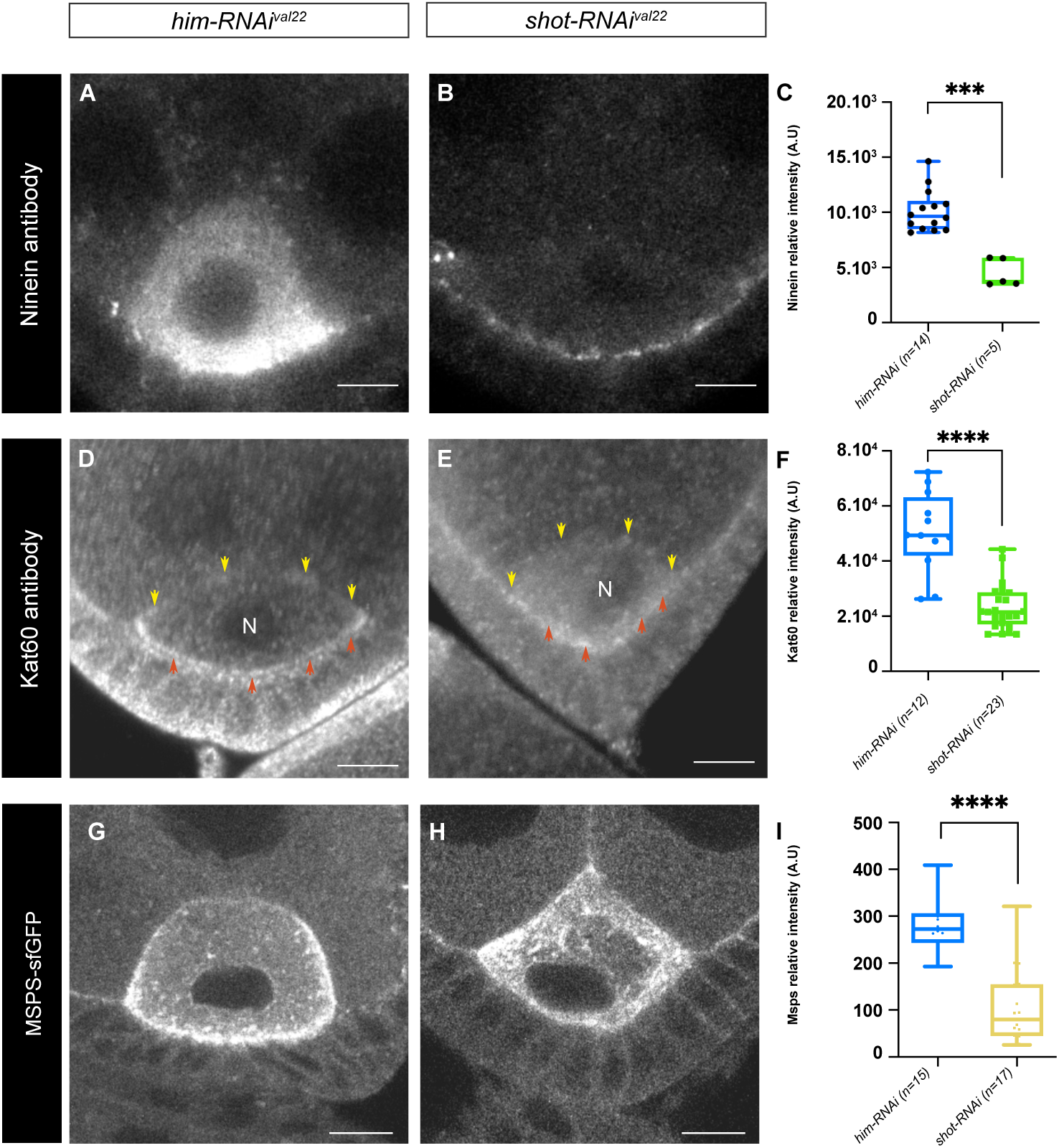
Shot functions as a scaffold for ncMTOC. **(A-C)** Representative fluorescent images illustrating that Ninein immunostaining intensity is strongly decreased at the posterior cortex of stage 6B *shot-RNAi^val^*^22^ oocytes (B) compared with *him-RNAi^val^*^22^ (A) oocytes used as controls. Quantification of signal intensity of Ninein immunostaining intensity at the posterior cortex (C). **(D-F)** Representative fluorescent images illustrating that Kat60 immunostaining intensity is strongly decreased at the posterior cortex of stage 6B *shot-RNAi^val^*^22^ oocytes (E) compared with *him-RNAi^val^*^22^ (D) oocytes used as controls. Quantification of signal intensity of Kat60 immunostaining intensity at the posterior cortex (F). **(G, I)** Representative fluorescent images illustrating that Msps-GFP intensity is strongly decreased at the posterior cortex of stage 6B *shot-RNAi^val^*^22^ oocytes (H) compared with *him-RNAi^val^*^22^ (G) oocytes used as controls. **(I)** Quantification of signal intensity at the posterior cortex of Msps-GFP intensity with *shot-RNAi^val^*^22^ oocytes compared with *him-RNA^val^*^22^. In all panels, the oocytes are oriented with anterior at the top and posterior at the bottom. Scale bars: 10µm. **** : p<0.0001; N: oocyte nucleus

## Discussion

Oocytes are highly specialized cells in which nuclear positioning is tightly regulated and must be precise to ensure successful fertilization. In many species, the nucleus must be off-centered to allow completion of meiosis and extrusion of polar bodies while minimizing cytoplasmic loss. However, an important preceding centering phase has been reported in several species, including zebrafish and mouse ^45–47^. In the latter, it has been shown that oocytes with prematurely off-centered nuclei are less prone to achieve correct meiotic maturation and that such oocytes are more frequently observed in old mice ^45^. In Zebrafish, the MT-actin crosslinking factor 1 (Macf-1), a member of the spectraplakin family as Shot, have been reported to be involve in nucleus centration, albeit in an atypical manner that is independent of MTs ^46^.

The coordination of nuclear positioning and movement relies on the cytoskeleton, involving either MTs or actin. This leads to another distinctive feature of oocytes: they lack centrosomes and therefore have evolved alternative MT-nucleation mechanisms that are independent of these canonical MTOCs.

We have identified a previously uncharacterized ncMTOC at the posterior plasma membrane of the *Drosophila* oocyte that assembles MTs during early oogenesis, prior to nuclear migration. This structure contributes to nuclear centering by generating MT-based pushing forces from the posterior cortex toward the oocyte center. Within this ncMTOC, Shot plays a central role by recruiting key MT regulators, including the severing enzyme Katanin, the polymerase Msps and minus-end associated protein Ninein, thereby promoting the formation of MTs that extend from the posterior cortex into the oocyte cytoplasm. MT organization at this stage also requires on γ-Tub37C function, independently of the centrosomes. γ-Tub37C is expressed in the germline and contributes to nuclear positioning as well as to MT density at the posterior cortex, yet it does not show specific accumulation at the oocyte cortex. This contrasts with canonical centrosomal proteins such as Asl and Sas-4, whose localization is restricted to centrosomes and whose depletion does not affect nuclear positioning or MT density. It also differs from Msps or Shot depletion, which strongly disrupt MT organization and impact oocyte size.

Altogether, these observations support the existence of a posterior ncMTOC that is at least partially independent of γ-Tub37C localization and suggest an additional, non-centrosomal role for γ-tubulin in MT organization within the oocyte.

Until now, the posterior cortex had not been considered a site with intrinsic MT-organizing potential, thus it could explain why in context where both centrosomes and Mud are depleted there are still cases where the nucleus still migrates compared to the situation in which oocytes are treated with a MT-depolymerazing drug (Colcemid or colchicine) ^7,10^. Thus, the identification of a posterior MTOC help unravel the complexity of the dense MT network of the oocyte. In addition, this discovery aligns with emerging evidence from other systems in which non-centrosomal MT formation occurs at specialized membrane domains, hinting at a conserved strategy for achieving spatially precise cytoskeletal control ^17,48^

The functional relevance of this posterior network is underscored by our direct readout of nuclear position, which provides an immediate mechanical report of MT-generated forces. This contrasts with several systems where outputs are more integrated or downstream, such as global cell polarity or synaptic vesicle localization in neurons. In those later cases, MT involvement is more indirect and intertwined with multiple roles of regulatory proteins. By relying on nuclear displacement, a proximal and mechanically interpretable metric, we directly link MT organization to force generation with minimal intervening processes. Thus, we will be able to further assess how different MT-networks integrate to control a common process, allowing at the same time a necessary redundancy for an important developmental process as well as a fine tuning of a balanced applied forces allowing fine tuning of nucleus positioning.

Our data also reveal that the formation or stabilization of this posterior acentrosomal network depends on Shot and Patronin, two proteins already known for their broad involvement in *Drosophila* oogenesis. Both proteins have been implicated in organizing MTs within the germline stem cell, thereby shaping the asymmetric arrays required for oocyte specification. Additionally, they contribute to MT organization at the ring canals involved in the nurse-cells to oocyte transport. The discovery that they also act at the posterior cortex further expands their functional repertoire and raises questions about how such versatility is achieved. A possible explanation lies in the extensive isoform diversity of both Shot (21 unique polypeptides) and Patronin (13 unique polypeptides), with different isoforms displaying distinct domain compositions and regulatory potentials. Isoform-specific interactions with cortical scaffolds, nucleators, or anchoring complexes could enable these proteins to support multiple, spatially segregated MT-organizing functions. Determining which isoforms operate at the posterior membrane and how they are differentially regulated will be essential to disentangle modular specialization from context-dependent deployment. For example, a specific isoform of Shot localizes to ring canals, where it co-localizes with actin filaments on the nurse cell side. It functions to regulate the directionality of cytoplasmic flow through these canals ^29^.

Recent studies have shown that at later stages (stage 9/10 and onwards), Shot is actively localized to, or maintained at, the anterior plasma membrane through Kinesin I activity ^49^. At stage 6, prior to nuclear migration, this mechanism alone is unlikely to account for the posterior enrichment of Shot, as MTs are thought to be predominantly oriented with their minus ends toward the posterior ^10,37^. Consistent with this, Khc loss-of-function mutants display nuclei predominantly at the anterior ^11^, opposite to the phenotype observed in *shot* mutants.

Interestingly, Shot is not strictly confined to the posterior cortex; low levels of Shot, Patronin, and other components are detectable at the anterior cortex. This suggests that multiple mechanisms may cooperate to establish Shot localization at early stages. We propose a model in which nuclear positioning at stage 6 is controlled by at least two partially overlapping pathways: (1) a Shot-dependent pathway, potentially involving subtle cortical cues or early MT dynamics, that biases Shot toward the posterior, and (2) a Kinesin-dependent mechanism that modulates centrosome-generated pushing forces, as previously described ^11^. In this framework, Shot enrichment at the posterior may result from the combined effects of these pathways, with relative contributions that shift as the oocyte develops.

This model highlights a dynamic, fine-tuned balance of forces and molecular localization, where early Shot recruitment sets the stage for later kinesin-dependent polarization. Future experiments will be needed to test how Shot is initially targeted and maintained at the cortex, how these pathways interact over time, and how they collectively integrate with MT architecture and cortical mechanics.

Although Actin has not previously been implicated in oocyte nuclear positioning, largely because of the limitations of available genetic and pharmacological tools and early requirements of Actin especially in oocyte specification, the finding that the Actin binding protein Shot may nonetheless suggest that actin may participate indirectly. It has been reported that Shot’s interactions are highly selective: it associates with specific actin structures rather than with all actin networks indiscriminately. This specificity raises the possibility that a specialized or previously overlooked actin population exists at the posterior cortex and contributes to the localized stabilization or anchoring of Shot and Patronin. Moreover, if Shot’s asymmetric localization depends on its ability to engage with such a distinct Actin structure, then the spatial patterning of this Actin network could itself dictate the polarized distribution and explain the dynamic relocalization of the Shot/Patronin complex, ultimately shaping the posterior acentrosomal MT array. Identifying the molecular identity, organization, and dynamics of this putative Actin population will therefore be a crucial direction for future studies.

ncMTOCs, which are based on the principles of MT anchoring, stabilization and nucleation, have been identified in multiple cell types and depend on a partially shared set of molecular components. Nevertheless, our findings, together with previous reports, suggest that the functional contribution of these components is highly context dependent. In the oocyte, ncMTOC nucleation activity appears to rely strongly on Msps, whereas in fat body cells Msps seems to primarily modulate MT stability ^14^. Similarly, the Katanin severing enzyme contribute to ncMTOC function in the oocyte but have been reported to be largely dispensable in fat body cells ^14^. Also, while individual depletion of Ninein and Patronin in fat body cells does not impair nuclear positioning, reflecting functional redundancy, there is no redundancy in the oocyte. Although differences in depletion efficiency or basal activity levels cannot be excluded, all these observations may also reflect distinct features of the underlying network organization. In fat body cells, the MT network is relatively stable and, as currently understood, relies on a single primary source at the nuclear envelope. In contrast, the oocyte MT network is more complex, involving at least three sources of microtubule polymerization, and must be dynamic to support directed nuclear migration, in a context where precise nuclear positioning is critical for proper subsequent embryonic development.

Severing enzymes, and Katanin in particular, are especially interesting, as precise regulation of their activity can produce different outcomes in the MT network. Accordingly, Katanin not only disassembles MTs but can also promote the formation of new ones by cutting long, stable polymers into shorter fragments that serve as seeds for further polymerization ^50^. This mechanism has been observed in *Drosophila*, both in larval sensory neurons and adult mechanosensory cilia ^42,43^.

Shot was identified as an interactor of Katanin 60 in a high-throughput yeast two-hybrid screen ^51^. Our results show that, in stage 6 oocytes, both Katanin 60 and Katanin 80 localize with Shot along the pPM. Importantly, this localization is lost upon Shot RNAi-mediated depletion.

Taken together, these findings suggest that Shot functions as a docking platform for Katanin-generated MT seeds, which are stabilized by MT minus-end–binding proteins such as Patronin and Ninein, and further elongated by the polymerizing factor Msps, thereby creating a microenvironment at the vicinity of the posterior plasma membrane that promotes and amplifies MT assembly.

## Materials and Methods

### *Drosophila* stocks and culture conditions

*Drosophila* stocks and crosses were maintained under standard conditions at 25°C. The following fly strains were used :

*w*^1118^ (BDSC#3605), *matTub-Gal4* (BDSC#7062), *Fs(2)Ket-GFP* ^52^, *ubi-PH-RFP* ^53^, *ubi-asl td-Tomato* gift of T. Avidor-Reiss ^54,55^, hsp-flp; FRT 82B pubi-GFP. pBac(Shot-YFP) ^56^, *matTub-Patronin-YFP* and CRISPR(*Patronin-YFP*) ^26^, *Kat80-YFP^CPTI^*^000764^ ^57^, obtained from the Kyoto *Drosophila* Stock Center, *γ-Tub23C-GFP* ^58^, *γ-Tub37C-mCherry* ^59^, *Grip75-sfGFP* ^59^*, Msps-sfGFP* (this study), *Sas-4^s^*^2214^ (Bloomington, BDSC#12119), *EB1::SSS::GFP* ^60^. Various information needed for this study was found in FlyBase ^61^

### RNAi crosses

The following fly lines, generated from the TRIP project ^62^ and obtained from the BDSC, have been used in this study:

*UAS-Him-RNAi^Val^*^22^ (attP2, Valium 22, GL01183) BDSC #42809, *UAS-CG12699-RNAi^Val^*^20^ (attP40, Valium20, HMS02833) BDSC #44111, *UAS-asl-RNAi^Val^*^20^ (attP2, Valium 20, HMS01453) BDSC #35039, *UAS-asl-RNAi^Val^*^22^ (attP40, Valium 22, GL00661) BDSC #38220, *UAS-shot-RNAi^Val^*^20^ (attP40, Valium 20, HMJ23381) BDSC#64041, *UAS-shot-RNAi^Val^*^22^ (attP2, Valium 22, GL01286) BDSC#41858, *UAS-Patronin-RNAi^Val^*^20^ (attP2, Valium20, HMS01547) BDSC #36659, *UAS-Kat60-RNAi^Val^*^22^ (attP40, Valium 22, GL00481) BDSC#35634, *UAS-kat-60L1-RNAi^Val^*^20^ (attP2, Valium 20, HMS00510) BDSC#32506, *UAS-kat80-RNAi^Val^*^20^ (attP40, Valium20, HMC06296) BDSC#66000, *UAS-γ-Tub37C-RNAi^Val^*^20^ (attP2, Valium 20, HMS00517) BDSC#32513, *UAS-grip75-RNAi^Val^*^20^ (attP40, Valium 20, HMS02789) BDSC#44072, *UAS-Sas4-RNAi^Val^*^20^ (attP2, Valium20, HMS01463) BDSC#35049, *UAS-msps-RNAi^Val^*^20^ (attP40, Valium20, HMS01906) BDSC#38990, *UAS-nin-RNAi^Val^*^20^ (attP40, Valium20, HMJ23837) BDSC#62414.

RNAi crosses were all performed using females from RNAi lines and maintained under standard conditions at 25°C. As control RNAi, we used lines expressing RNAi directed against genes not expressed in ovary, i.e CG12699 and Holes in Muscle (Him) ^63,64^. In addition, the lines were selected and used regarding the Valium plasmid used as well as the insertion point in the genome (attP2 or attP40).

### Generating endogenously tagged Msps-sfGFP fly lines

Msps-sfGFP endogenously tagged lines were generated using CRISPR-mediated homologous recombination. A homology-repair vector was used containing: 1100 bp upstream homology arm -GGS(X5) linker-sfGFP – 1024 bp downstream homology arm - LoxP-mini-white-LoxP cassette. Homology vectors were constructed via HiFi assembly (NEB) using PCR fragments from the following templates: LJ46 (backbone, gift from the Emmanuel Derivery lab) for the LoxP-White-LoxP cassette; genomic DNA prepared from *nos-Cas9* flies (using MicroLYSIS, Clent Life Science) for the upstream and downstream homology arms; and plasmid 1314 (DGRC) containing the sfGFP tag. The following guide RNA sequence was used: ACGGGAAGCGCACAGTTTAT. This sequence was cloned into the pCFD3 vector as described in ^65^. The homology and guide vectors were co-injected into the *nos-Cas9* fly line (*yw; nos-Cas9(II-attP40)*) by the BestGene facility. Injected flies were screened using the mini-white selection marker and then the selection cassette was excised by crossing to a line containing Cre-recombinase (BDSC #851), then verified by PCR and genomic DNA sequencing. After balancing, healthy homozygous stocks were established and maintained.

### Heat-Shock Treatment for clonal analysis

Heat-shocks were carried out for 1 hr in a water bath at 37°C, 3 days in a row from L1 larvae.

### Immunostaining of the fly ovaries

1day old females were collected and put with fresh media for 30-48 hours at 25°C prior to dissection. Ovaries were dissected, fixed in PBS with 4% paraformaldehyde and incubated overnight at 4°C with primary antibodies in PBS with 0,1% Tween. Primary antibodies include: mouse α-Shot (DSHB #mAbRod1, supernatant) at 1:10; mouse α-Kat60 (gift from S. Rogers ^66^, supernatant) at 1:2000; rabbit α-Ninein (from ^67^) at 1:500; rabbit Anti-Msps (from ^68^) at 1:200. Ovaries were then incubated with secondary antibodies at room temperature for 2 hours. Secondary antibodies include: chicken α-mouse Alexa647 (Invitrogen, #A21463) at 1:200 and goat α-rabbit Alexa647 (Jackson Immunoresearch #111-605-144) at 1:200. For egg chambers that did not express a plasma membrane marker nor a nucleus marker, ovaries were respectively incubated with Alexa Fluor 488 Phalloidin (Invitrogen, #A12379) at 1:100 and with Wheat Germ Agglutinin (WGA) (Molecular Probes) at 1:200, at 4°C overnight.

After washes, ovaries were mounted in Citifluor^TM^ (EMS). Images were captured using Zen software on a Zeiss 710 confocal microscope (488nm, 561nm, 640 nm lasers and the x40 objective).

### Visualization of the oocyte MT network

One-day-old females were collected and transferred onto fresh food medium for 24–48 hours at 25°C prior to dissection. Ovaries were dissected in Schneider’s *Drosophila* medium and immediately incubated with SiR-Tubulin (Spirochrome) diluted 1:200 in Schneider’s medium for 30 minutes at room temperature with gentle agitation. After incubation, ovaries were mounted in halocarbon oil (Voltalef 10S) on coverslips for imaging.

Images were captured using Metamorph software on a Zeiss Axio Observer Z1 confocal microscope coupled with a spinning disk module CSU-X1 and a sCMOS camera PRIME 95 (488nm, 561nm, 640 nm lasers and a x63 oil immersion objective). In order to image the whole nucleus, 21 stacks along the z axis of a 1µm range were taken for each egg chamber.

Quantification at the posterior cortex: Signal quantification was performed at the posterior cortex using a manually defined region of interest (ROI). The ROI consisted of a 10-pixel-wide (1px = 0,091 µm) line drawn along the posterior cortical region within the oocyte, and the integrated fluorescence intensity within this region was measured. To account for inter-sample variability in signal intensity and imaging conditions, a normalization procedure was applied using adjacent follicular cells that are not affected by the expressed RNAi as an internal reference. For each sample, three lateral follicular cells in direct contact with the oocyte were selected, and their fluorescence intensity was quantified. A correction factor was calculated by dividing the maximum signal intensity measured in the sample by the mean signal intensity of the selected follicular cells. This correction factor was subsequently applied to the total fluorescence intensity measured within the posterior cortical ROI, yielding normalized intensity values for comparative analysis across samples.

### Image analysis, quantification and statistical analysis

All images were processed using the software Fiji ^69^

#### Egg chamber staging

To stage the egg chambers, we proceeded as described in ^11^. In brief, we measured nucleus diameters of the two closest nurse cells of the oocyte, using the z-section corresponding to the larger diameter of the considered nurse cells. These diameters were measured using the « Straight, segmented lines » tool on Fiji. In case the two nurse cells show diameters that can be categorized in different stages, we then took in account the shape and the size of the oocyte and the egg chamber aspect ratio as suggested by ^70^.

#### Determination of nuclear position in the oocyte

Nucleus position was mainly assessed on a qualitative criterion, nevertheless in case of uncertainty, we have applied a quantitative method consisting in making the ratio of the distances: hemisphere anterior of the nucleus - anterior membrane (d_ant_) and hemisphere posterior of the nucleus - posterior membrane (d_post_); r = d_ant_/d_post_. If r < 0,5 the nucleus was categorized as “anterior”, if r > 2 the nucleus is determined as “posterior” and if 2 > r > 0,5, the nucleus is considered as “central”.

#### Statistics

Bar plots as well as statistical tests were carried out with GraphPad Prism10.

### Live imaging of MT dynamics in *Drosophila* oocytes

One-day-old transgenic females expressing EB1::SSS::GFP were collected and transferred onto fresh food medium for 24–30 hours at 25°C prior to dissection. Ovaries were dissected in Schneider’s *Drosophila* and mounted in halocarbon oil (Voltalef 10S) on coverslips for live imaging.

Images were acquired using Metamorph software on a Zeiss Axio Observer Z1 microscope equipped with a CSU-X1 spinning disk module and a PRIME 95B sCMOS camera, using a 488 nm laser and a 100× oil immersion objective (NA 1.4). Time-lapse series were recorded on a single focal plane at 500 ms intervals for 2 minutes for each egg chamber to monitor MT growth dynamics.

### Image Processing and EB1 Comet Tracking

#### Image Processing

Time-lapse image stacks were processed using Fiji. First, the stacks were reoriented such that the oocyte was positioned at the bottom of the field of view. Regions of interest (ROIs) were manually defined to delineate the oocyte boundary, as well as a 10-pixel-wide (1px = 0,091 µm) ROI along the posterior plasma membrane. These ROIs were combined into a three-channel hyperstack (oriented movie, posterior ROI, and oocyte) suitable for subsequent analysis in IMARIS software (Bitplane).

#### Comet Segmentation and Tracking

EB1 comets were automatically tracked using the IMARIS tracking module. The segmentation parameters were defined by specifying the approximate comet diameter (0.3 µm) and a quality threshold based on the signal-to-noise ratio to ensure accurate detection. For comet tracking, only comets that persisted for more than 2.5 s in the imaging plane (corresponding to at least five consecutive detected objects) were considered. This criterion excluded trajectories that left the focal plane along the z-axis.

To minimize the merging of trajectories arising from distinct comets in the dense MT network, a trajectory was terminated if a comet disappeared for a single time point (0.5 s). Additionally, a maximum distance of 1 µm between consecutive trajectory points was set, corresponding to the maximum average distance traveled by a comet in a single time frame.

Tracks that included at least one time point within the posterior ROI were classified as posterior trajectories.

#### Trajectory Analysis and Categorization

Trajectory vectorization was performed using a custom R script, which calculated the vector from the starting point to the end point of each track. The angle of this vector relative to the horizontal axis was then determined. Based on the orientation of the oocyte, comets emerging from the posterior plasma membrane exhibited negative angles and were categorized as posterior Plasma Membrane Arising Comets (pPMACs).

#### Statistics

Bar plots as well as statistical tests were carried out with GraphPad Prism10.

## Acknowledgements

We would like to thank Nicolas Destombes, Lois Goasmat-Arnold and Cassandra Dubois for preliminary functional analyses. We thank the ImagoSeine core facility of the Institut Jacques Monod, member of IBiSA and the France-BioImaging (ANR-10-INBS-04) infrastructure at the Institut Jacques Monod for their help and support. Dmitry Nashchekin, Katja Röper, Stephen Rogers, Emmanuel Derivery FlyBase release (e.g. FB2022_03) used for data obtained and analyzed using FlyBase, the Bloomington *Drosophila* Stock Center, Vienna *Drosophila* Research Center and the Developmental Studies Hybridoma Bank for fly stocks and reagents. We are grateful to Guichet Lab members for helpful discussions during the course of this study and their comments on the manuscript.

## Competing interests

The authors declare no competing or financial interests.

## Author contributions

The project was conceived by F. Bernard and A. Guichet. F. Bernard. and A. Guichet designed the experiments that were subsequently performed by F. Roland-Gosselin, S. Sahayan Nevil Fernando and F. Bernard. M. Peroche wrote the R script necessary for the EB1 comet analysis. The data were analyzed by F. Roland-Gosselin, F. Bernard and A. Guichet. The Msps-sfGFP line was generated by A. Yagoubat and P. Conduit. The project funding, administration, and supervision were provided by F. Bernard and A. Guichet. F. Roland-Gosselin prepared the figures. A. Guichet and F. Bernard wrote the manuscript, which was edited and reviewed by all authors.

## Fundings

This work was funded by the Fondation ARC (PJA-20181208148) pour la Recherche sur le Cancer, the Ligue Contre le Cancer Comité de Paris (RS20/75-17), the Association des Entreprises contre le Cancer (Grant Gefluc 2020 #221366), an Emergence grant from IdEx Université de Paris (ANR-18-IDEX-0001) and by ANR grant NACFLO. F.R-G was supported by a fellowship from “Ministère de l’Education Nationale, de la Recherche et de la Technologie” (MENRT) obtained from the BioSPC doctoral school and ARC fellowship. F.B is supported by Université Paris Cité. A.G is supported by the CNRS.

## Notes

### Competing Interest Statement

The authors have declared no competing interest.

