## Supplementary material for "The Spectraplakin Short Stop (Shot) Organizes an Acentrosomal Microtubule Network in Early Oogenesis, Essential for Nuclear Positioning": Sup Fig 1

### Fig. sup. 1

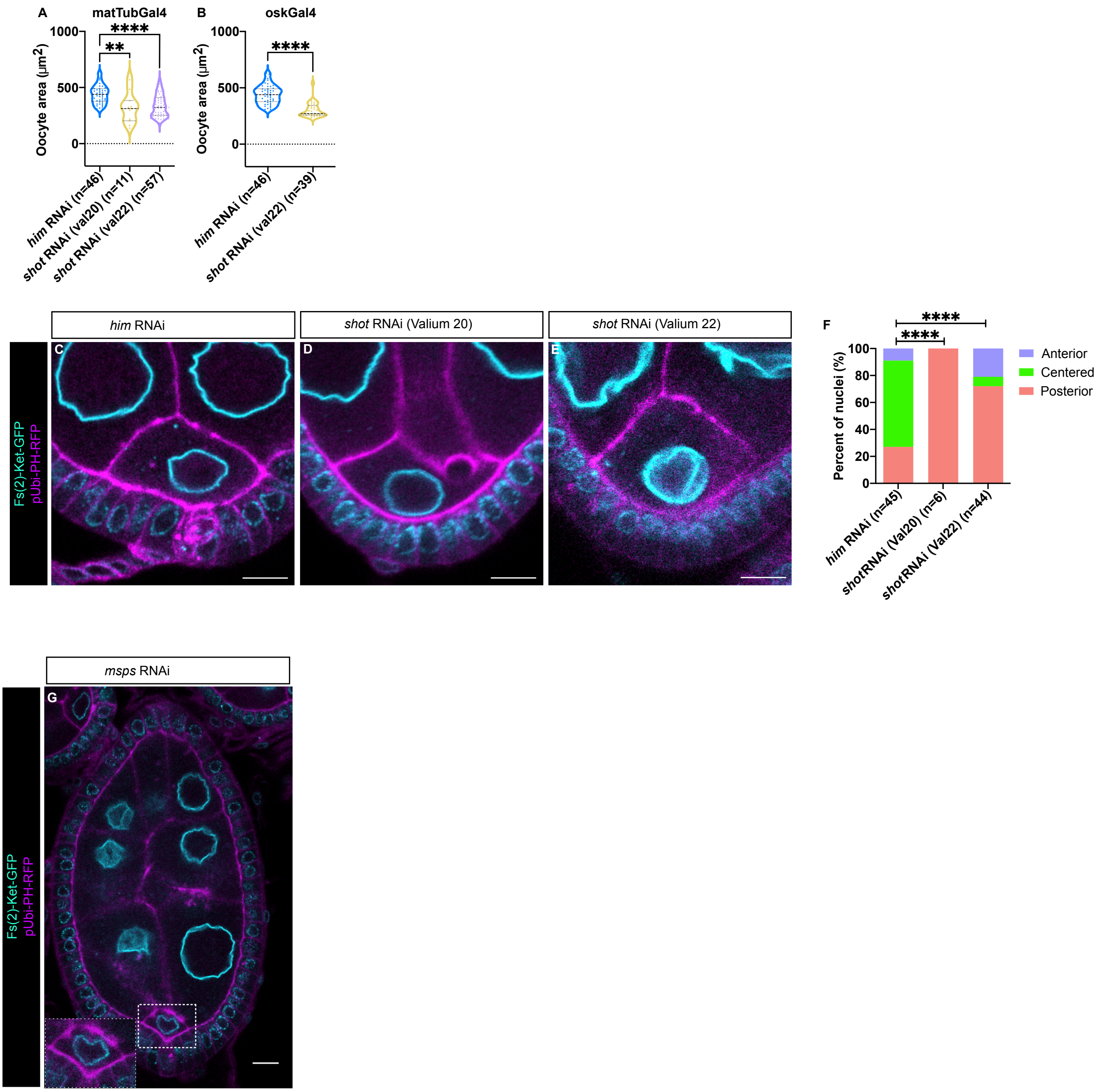


**Supplementary Figure 1:**
**(A)** Quantification of the oocyte area for *Him-RNAi* as control, *shot-RNAi*$^{val\ 20}$ and *shot-RNAi*$^{val\ 22}$ conditions under the control of *matTub-Gal4* transgene. *shot-RNAi*$^{val\ 20}$ and *shot-RNAi*$^{val\ 22}$ are two independent RNAi lines. **(B)** Quantification of the oocyte area for *Him-RNAi* as control and *shot-RNAi*$^{val\ 22}$ conditions under the control of *oskar-Gal4* transgene. **(C-E)** Representative images illustrating the position of the nucleus in stage 6B oocyte for *Him-RNAi* as control (C), *shot-RNAi*$^{val20}$ (D) and *shot-RNAi*$^{val\ 22}$ (E) conditions. The nuclear envelope (cyan) is highlighted by Fs(2)Ket-GFP and the plasma membrane (magenta) by PH-RFP. **(F)** Quantification of the proportions of nuclear positions in the control, *shot-RNAi*$^{val\ 20}$ and *shot-RNAi*$^{val\ 22}$ groups. Positions have been categorized and color-coded as anterior in pale blue, center in green, and posterior in red. n indicates the number of analyzed egg chambers. **(G)** Representative image illustrating oocyte growth defects induced with msps-RNAi. In all panels, the oocytes are oriented with anterior at the top and posterior at the bottom. Scale bars: 10µm. * : p<0.05 ; ** : p<0.01 ; *** : p<0.001 ; **** : p<0.0001; ns, not significant. N: oocyte nucleus.
